# Diverse RNA viruses associated with diatom, eustigmatophyte, dinoflagellate and rhodophyte microalgae cultures

**DOI:** 10.1101/2022.05.14.491972

**Authors:** Justine Charon, Tim Kahlke, Michaela E. Larsson, Raffaela Abbriano, Audrey Commault, Joel Burke, Peter Ralph, Edward C. Holmes

## Abstract

Unicellular microalgae are of immense ecological importance with growing commercial potential in industries such as renewable energy, food and pharmacology. Viral infections can have a profound impact on the growth and evolution of their hosts. However, very little is known of the diversity within, and effect of, unicellular microalgal RNA viruses. In addition, identifying RNA viruses in these organisms that could have originated more than a billion years ago constitutes a robust data set to dissect molecular events and address fundamental questions on virus evolution. We assessed the diversity of RNA viruses in eight microalgal cultures including representatives from the diatom, eustigmatophyte, dinoflagellate, red algae and euglenid groups. Using metatranscriptomic sequencing combined with bioinformatic approaches optimised to detect highly divergent RNA viruses, we identified ten RNA virus sequences, with nine constituting new viral species. Most of the newly identified RNA viruses belonged to the double-stranded *Totiviridae, Endornaviridae* and *Partitiviridae*, greatly expanding the reported host range for these families. Two new species belonging to the single-stranded RNA viral clade *Marnaviridae*, commonly associated with microalgal hosts, were also identified. This study highlights that a great diversity of RNA viruses likely exists undetected within the unicellular microalgae. It also highlights the necessity for RNA viral characterisation and to investigate the effects of viral infections on microalgal physiology, biology and growth, considering their environmental and industrial roles.

**Importance:** In comparison to animals or plants, our knowledge of the diversity of RNA viruses infecting microbial algae – the microalgae – is minimal. Yet describing the RNA viruses infecting these organisms is of primary importance at both the ecological and economical levels because of the fundamental roles these organisms play in aquatic environments and their growing value across a range of industrial fields. Using metatranscriptomic sequencing we aimed to reveal the RNA viruses present in cultures of eight microalgae species belonging to the diatom, dinoflagellate, eustigmatophyte, rhodophyte and euglena major clades of algae. This work identified ten new divergent RNA virus species, belonging to RNA virus families as diverse as the double-stranded *Totiviridae, Endornaviridae, Partitiviridae* and the single-stranded *Marnaviridae*. By expanding the known diversity of RNA viruses infecting unicellular eukaryotes, this study contributes to a better understanding of the early evolution of the virosphere and will inform the use of microalgae in industrial applications.

## Introduction

Viruses are often considered the most ancient “life forms’’ (*i*.*e*. replicatory agents). As studies of the viromes of increasingly diverse taxa proceed, the more their remarkable ubiquity, diversity and abundance becomes apparent^1^. RNA viruses are by far the most abundant microorganisms in marine systems^2^ and play fundamental roles in these environments by infecting and regulating phytoplankton populations^2^. RNA viruses that infect unicellular photosynthetic microalgae are also of primary importance for marine resource management due to the significant ecotoxicological effect of some microalgal hosts, including abundant dinoflagellate species^3^. There is also growing awareness of the value of microalgal cultures for biofuels, pharmacology, water treatment, food and the aquacultural industries^4–7^. Indeed, the intensive commercial cultivation and production of microalgae populations could be seriously impacted by viral disease outbreaks^8^. Accordingly, an extensive description of the RNA virus diversity in unicellular microalgae is of importance to better understand their role and impact on natural microalgal populations and in anticipating the consequences of industrial cultivation.

Knowledge of the RNA virosphere in overlooked eukaryotic lineages that evolved billions of years ago – such as the microalgae – could significantly enhance our understanding of the earliest events in RNA virus evolution. With barely 100 species of RNA viruses reported since the first isolation of a microalgae-infecting RNA virus in 2003^9^, our current knowledge of RNA viruses infecting microalgae is limited, representing less than 0.5% of the RNA viruses for which hosts have been formally established^10^. This lack of knowledge most likely reflects the historical focus on viruses that cause disease in humans and bioresources (domestic animals, animal and insect vector, plants) rather than those infecting microbial eukaryotes.

The study of global viromes has been revolutionised by metagenomics. By avoiding cultivation limitations and paving the way for the exploration of very diverse environments (soil, water, etc.), the metagenomic era has multiplied the number of RNA viruses described by many thousands^10–13^. This is particularly evident in the field of “phycovirology” (the study of algal viruses), for which recent studies investigating RNA viruses using metagenomic approaches have revealed a high diversity and prevalence of RNA viruses in several microalgae lineages^11,14–21^. While the positive-sense single-strand (ss+) picorna-like *Marnaviridae* are the best described family of microalgae-infecting viruses^11,22,23^, metagenomic studies continue to expand the diversity of microalgal viruses, including identification of the double-strand (ds) RNA viruses from the orders *Ghabrivirales* (*Totiviridae*-like), *Durnavirales* (*Partitiviridae*-like) and *Martellivirales* (*Endornaviridae*-like)^19,20,24–26^, as well as ss+ RNA viruses from the *Sobelivirales* (*Alvernaviridae*), *Nodamuvirales* (*Nodaviridae*), *Wolframvirales* (*Narnaviridae*) and *Cryppavirales* (*Mitoviridae*) phyla^19,20,24,27,28^. To date, the majority of the microalgal hosts documented to contain RNA viruses are from the Bacillariophyta (diatom) and Dinoflagellata (dinoflagellate) lineages. However, some viruses have been reported from other stramenopile hosts (such as Phaeophytes, Raphidophytes and Xanthophytes)^9,20,28–30^ and in some other major groups of microalgae such as the *Rhizaria* ^9,20^, Chlorophyta^19,31^, Rhodophyta^20,25,26^ and more recently Haptophyta^20^.

To increase our understanding of the RNA virosphere in microalgae we assessed the diversity of RNA viruses in eight microalgal species, covering the major groups of stramenopiles, including eustigmatophytes (*Nannochloropsis oceanica, Nannochloropsis oculata)* and diatoms *(Thalassiosira weissflogii*), alveolates including dinoflagellates (*Prorocentrum* cf. *balticum, Prorocentrum lima, Gambierdiscus carpenteri*), red algae (*Rhodella maculata*) and euglenid (*Euglena gracilis*). By using a “culture-based” metatranscriptomic approach we combined the power of unbiased detection of ultra-large-scale RNA sequencing with the use of mono-organism culture to assist in associating the viruses identified to their specific algae hosts. Given the high levels of sequence diversity observed in many RNA viruses, we paid particular attention to identifying divergent virus-like sequences.

## Results and Discussion

We searched for RNA virus sequences associated with cultures of eight unicellular microalgal species, representing four major algal groups: stramenopiles, alveolates, rhodophytes and euglenozoa (Figure 1).

**Figure 1.**
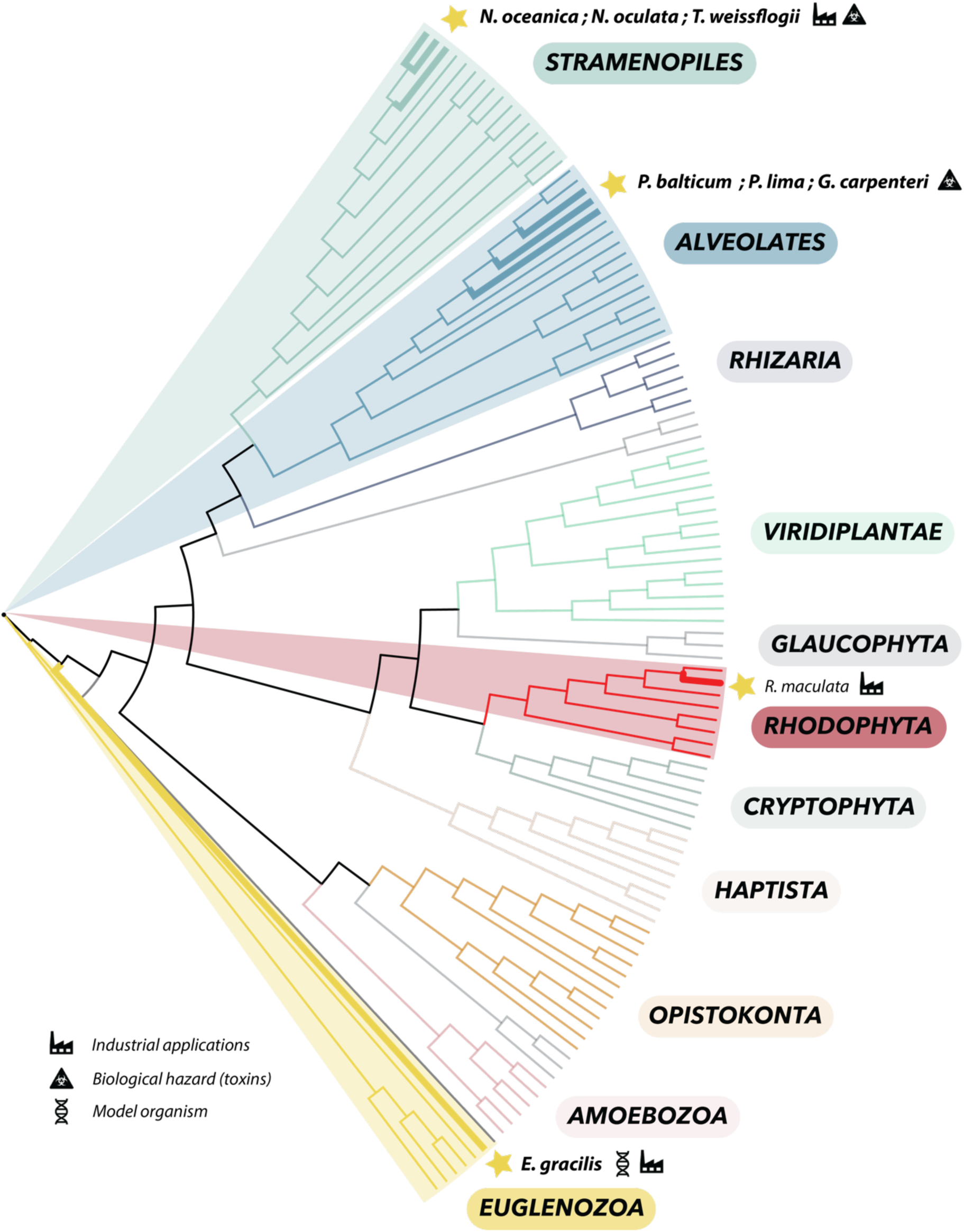
Phylogenetic position of microalgal species used in this study among the global Eukaryote phylogeny. Microalgal species names are indicated in italics and their main applications (industrial, biological hazard (toxins) and model organisms) are specified by icons. Species for which no RNA viruses were reported prior to this study are indicated in bold. The Eukaryote cladogram was adapted from ref.^32^.

Following total RNA extraction from each microalgal culture, metatranscriptomic sequencing was used to obtain deep transcriptomes. The corresponding RNAs and data yields for each microalgal sample/library are detailed in Table A1 and Figure A1. By combining a standard metagenomic bioinformatic pipeline with the protein HMM-profile and structural comparison developed in RdRp-scan^33^ we were able to identify ten new viral-like sequences (Table 1). With the exception of the unicellular red algae *Rhodella maculata*, recently associated with the *Despoena mito-like virus*^20^, these represent the first reports of viruses in each microalgal species investigated (Figure 1). The ten viral sequences found in this study were compared to the genomic sequences of the corresponding algal host whenever possible (Table A2). Accordingly, nine of the ten viral sequences identified were not found in the host genome and therefore treated as exogenous viruses (Table 1). In contrast, the viral signal detected from *Euglena gracilis* using HMM-based approach was identical to *Euglena* genome sequences (Table 1) and hence likely corresponds to an endogenous viral element (EVE; see below).

**Table 1.** List of new viruses and endogenous viral elements found in this study. EVE: endogenous viral element. Full-length versus partial information was hypothesized from genomic length and organisation.

| Virus name | Host (lineage) | RdRp phylum/order related | RT-PCR validated?<br>Virus/Host | Coverage<br>(quality/nb reads) | length | Full-length vs partial | Exogenous vs EVE |
| --- | --- | --- | --- | --- | --- | --- | --- |
| Taphios ghabri-like virus 1 | *Nannochloropsis oceanica*<br>(Eustigmatophyte) | Duplornaviricota/Ghabrivirales | YES/YES | GOOD/13,769 | 4876 | Likely Complete | Exogenous |
| Taphios ghabri-like virus 1 | *Thalassiosira weissflogii*<br>(Eustigmatophyte) | Duplornaviricota/Ghabrivirales | YES/YES | GOOD/876 | 4835 | Likely Complete | Exogenous |
| Triopas ghabri-like virus 1 | *Prorocentrum cf. balticum*<br>(Dinophyceae) | Duplornaviricota/Ghabrivirales | NO/YES | AVERAGE/70 | 1425 | Partial | Exogenous |
| Diktys durna-like virus 1 | *Prorocentrum lima*<br>(Dinophyceae) | Pisuviricota/Durnavirales – Partitivirus | YES/YES | GOOD/24,826 | 1826 | Partial (1 segment missing ?) | Exogenous |
| Orion durna-like virus 1 | *Gambierdiscus carpenteri*<br>(Dinophyceae) | Pisuviricota/Durnavirales | YES/YES | GOOD/1,198 | 2077 | Partial (1 segment missing ?) | Exogenous |
| Almopos endorna-like virus 1 | *Gambierdiscus carpenteri*<br>(Dinophyceae) | Kitrinoviricota/Martellivirales | YES/YES | GOOD/54,823 | 21494 | Likely Complete | Exogenous |
| Althepos endorna-like virus 1 | *Gambierdiscus carpenteri*<br>(Dinophyceae) | Kitrinoviricota/Martellivirales | YES/YES | GOOD/460 | 4825 | Likely Complete | Exogenous |
| Phineus pisuviri-like virus 1 | *Rhodella maculata*<br>(Rhodophyta) | Pisuviricota/Picornavirales | YES/YES | GOOD/14,972 | 6398 | Likely Complete | Exogenous |
| Megareus marna-like virus 1 | *Nannochloropsis oculata*<br>(Stramenopiles) | Pisuviricota/Picornavirales | NO/NO | GOOD/425 | 1222 | Partial | Exogenous |
| Minyas marna-like virus 1 | *Nannochloropsis oculata*<br>(Stramenopiles) | Pisuviricota/Picornavirales | NO/NO | GOOD/472 | 1130 | Partial | Exogenous |
| Pisuviri-like signal | *Euglena gracilis*<br>(Euglenozoa) | Pisuviricota/Uncertain placement | YES/YES | GOOD/1,531 | 1484 | EVE | EVE |

To eliminate contamination during the library preparation or sequencing, we tested the presence of all viruses in total RNA samples using RT-PCR (Figure A2). This resulted in the detection of 8 of the 11 virus/samples tested (Table 1 and Figure A2). Triopas ghabri-like virus 1 could not be detected in *P*. cf. *balticum* RNAs (Figure A2), likely because of the very low abundance of this viral contig (Table 1). Megareus marna-like virus 1 and Minyas marna-like virus 1, both associated with the *N. oculata* sample, similarly could not be confirmed using RT-PCR. In addition, the positive control used to target the *N. oculata* internal transcribed spacer (ITS) sequence did not return any PCR signal (Figure A2). Hence, the meagre quantity of total RNA extracted from *N. oculata* cultures may explain the difficulty in validating both host gene and associated viruses using RT-PCR (Table A1).

### Placement of the newly identified microalgal viruses within global RNA virus diversity

To characterise the newly-identified viruses, we first used phylogenetic analysis to place the new viral sequences within the diversity of viral RNA-dependent RNA polymerase (RdRp) sequences at the phylum level using the recently developed RdRp-scan resource^33^. These large-scale phylogenies show that the viral sequences identified fell in diverse positions among those RNA viruses identified to date, with two belonging to the *Duplornaviricota*, two falling into the *Kitrinoviricota*, and six sharing homologies at amino acid level with *Pisuviricota* viruses (Figure 2).

**Figure 2.**
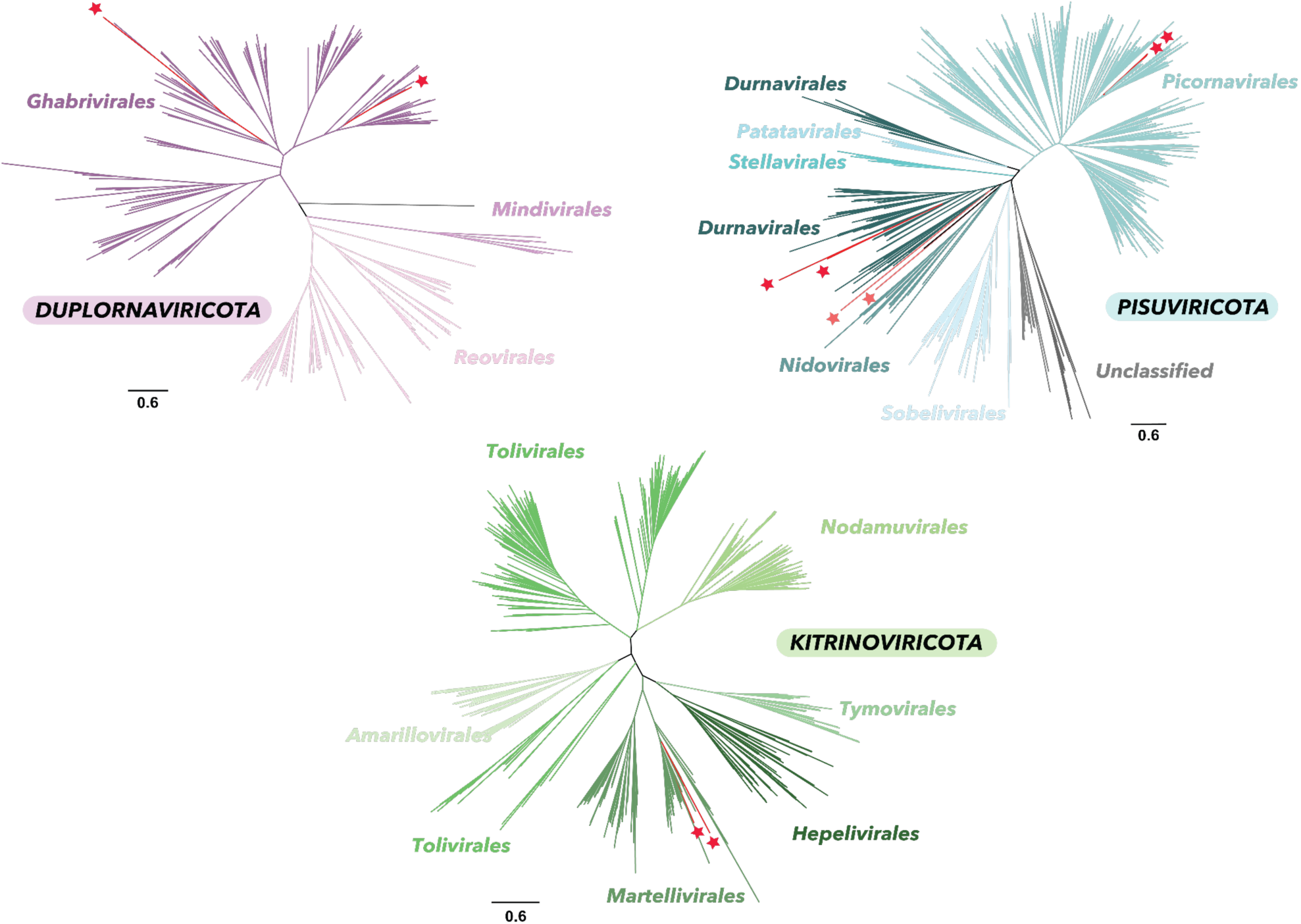
Phylogenetic placement of the newly identified viruses within the diversity of *Riboviria* phyla. Unrooted ML trees. Red stars indicate the viruses newly identified. Light red stars represent RdRp-like sequences obtained using the HMM-based RdRp-scan method. Scale bars represent the number of amino acid substitutions per site.

We then conducted additional phylogenetic analyses focusing on the viral sub-clades that contained the ten newly identified sequences. These comprised the *Ghabrivirales, Endornaviridae, Durnavirales* and *Marnaviridae* lineages and are described below.

### New microalgae-infecting viruses suggest a TSAR-infecting Totiviridae genus

Among the ten viral sequences identified in this study, two were related to *Totiviridae*-like viruses (Table 1 and Figure 2). Triopas ghabri-like virus 1, identified in the dinoflagellate *P*. cf. *balticum*, forms a clade with Arion toti-like virus identified in the dinoflagellate *P. bahamense*^20^. Together, these two viruses group with those previously reported in microalgae and oomycete hosts (Figure 3).

**Figure 3.**
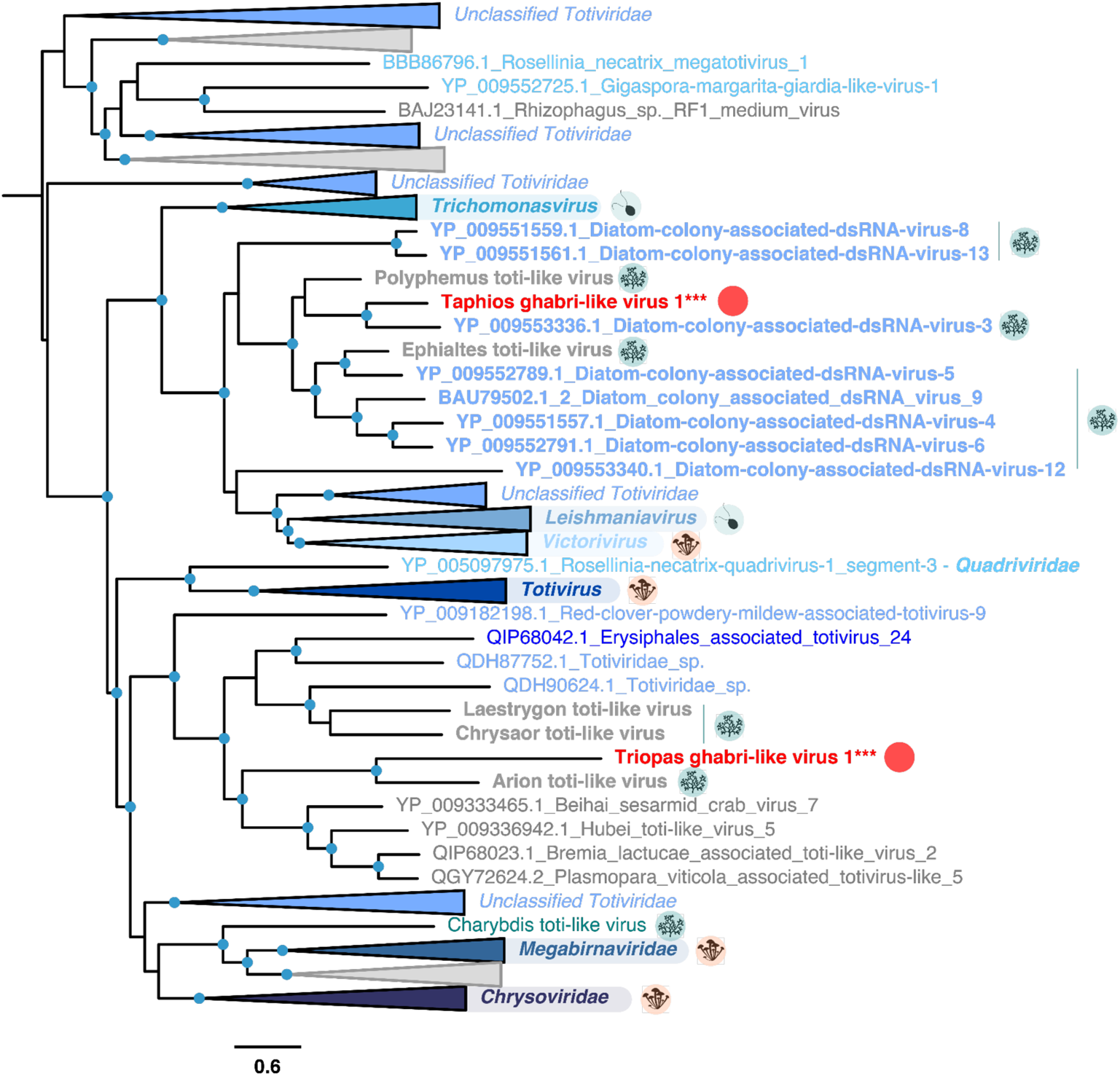
Phylogeny of the *Ghabrivirales*. Sequences in grey denote previously unclassified viruses while those in bold refer to microalgae-associated viruses. Host lineages are indicated in circles to the right of major viral clade labels and correspond to fungi (orange), protozoa (light blue) and microalgae (blue). The new viral sequences identified in this study are indicated with a red circle. The tree is mid-rooted and confident nodes (with SH-alrt likelihood ratio test values >=80%) are represented as circles. The scale bar represents the number of amino acid substitutions per site.

The short length of the RdRp-encoding segment identified here – 1.4 kb – suggests that the genome of *Triopas ghabri-like virus 1* is partial (Figure A3). While this limits the discussion of genomic attributes, an additional open reading frame (ORF) in reverse-orientation and without any known function associated, was predicted using the standard genetic code. Such use of anti-sense ORF would constitute an original feature in the *Totiviridae* (Figure A3). Triopas ghabri-like virus 1 associated RNAs were found at very low abundance in the *P*. cf. *balticum* sample and could not be confirmed experimentally by RT-PCR (Table 1, Figure A2). Although this viral sequence requires additional validation, it supports previous suggestions of dinoflagellate-infecting *Totiviridae*^20^ and constitutes further evidence for recognizing a new TSAR-infecting (Telonemid, Stramenopile, Alveolate and *Rhizaria* supergroup) genus within the *Totiviridae*^20^.

A second Toti-like virus, Taphios ghabri-like virus 1, was found in the eustigmatophyte *N. oceanica* and the diatom *T. weissflogii*. It forms a clade with the algae-associated Polyphemus and Ephialtes toti-like viruses, both previously identified in *Astrosyne radiata* (diatom) samples^20^. They also form a sister clade to the *Trichomonasvirus, Victorivirus* and *Leishmaniavirus* genera, infecting protozoan parasites and fungi^34–36^. To consolidate the host-virus relationship and potentially elongate the genomic sequence, we screened for the presence of the newly described viruses in additional host transcriptomes available in the Sequence Read Archive (SRA) (Table A3). Accordingly, Taphios ghabri-like virus 1 sequences were observed in one transcriptome (SRR12347810) of the diatom *Phaeodactylum tricornutum*, with only eight single nucleotide polymorphisms (SNP) reported at the genome level.

The total length of the Taphios ghabri-like virus 1 sequence, at 4.8kb, is in the range of other *Totiviridae* and, along with the read coverage profile, suggests that the full-length genome has been obtained (Figure A3). The organisation of the Taphios ghabri-like virus 1 genome into two overlapping ORFs, probably translated with a +1 ribosomal frameshift, corresponds to the genomic features commonly observed among the *Totiviridae*. The first ORF likely encodes a coat protein, while no annotations could be retrieved from InterProscan analysis for this ORF^37^. We hypothesise from the placement within the *Totiviridae* phylogeny and the similarities in genome organisation and length that this virus has a dsRNA genome. Combined, the results from RdRp phylogenies, genome organisation and host range are in accord with establishing a new *Totiviridae* genus infecting diatom and eustigmatophyte hosts.

The observation of *Totiviridae* likely infecting dinoflagellates, diatoms and eustigmatophyte hosts aligns with the suspected ubiquity of these dsRNA viruses in microalgae^20^ and unicellular eukaryotes more generally^38^. Notably, the *Totiviridae* have been associated with changes in host fitness and to hyper- or hypovirulence of some of their hosts^39–41^. The effects of the newly discovered *Totiviridae* genus on corresponding dinoflagellate, diatom and eustigmatophyte microalgal cultures require additional investigation and could be of interest considering their potential effects on growth, including that of harmful algal blooms (HABs), and commercial cultivation yields.

### First association of alphaendornavirus with dinoflagellates

Two of the viruses identified in this study cluster within the *Endornaviridae* family of dsRNA viruses. Specifically, Althepos endorna-like virus 1 and Almopos endorna-like virus 1 (both retrieved from *Gambierdiscus carpenteri* – Dinophyceae) group with members of the alphaendornavirus genus, a genus within the *Endornaviridae* previously associated with land plants, fungi and oomycetes^42^ (Figure 4).

**Figure 4.**
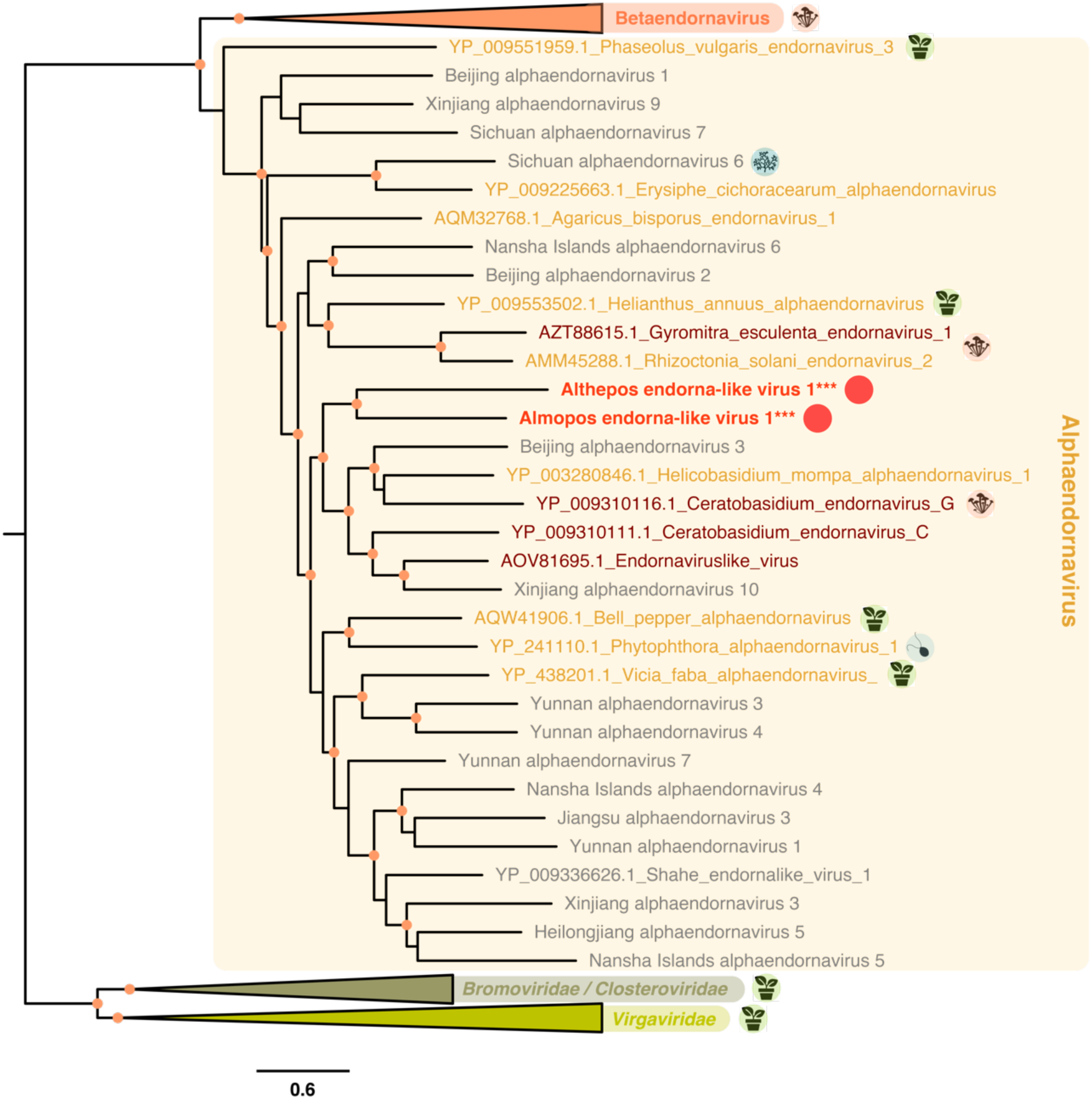
Phylogeny of the *Endornaviridae*. Sequences in grey indicate unclassified viruses and sequences in bold refer to microalgal-associated viruses. Host lineages are indicated in circles to the right of major viral clades and correspond to fungi (orange), land plants (green), protozoa (light blue) and microalgae (blue). The new viral sequences identified in this study are indicated with a red circle. The Sichuan alphaendornavirus cluster includes the diatom-associated RNA virus 15, previously reported from diatom-containing samples^18^. The tree is mid-rooted and confident nodes (with SH-alrt likelihood ratio test values >=80%) are represented as orange circles. The scale bar represents the number of amino acid substitutions per site.

*Endornaviridae* dsRNA genomes are 9.7 to 17.6 kb in length and encode a single polyprotein with a RdRp domain located in the Cter^38^. The genome organisation of Almopos endorna-like virus 1 therefore possesses features common to the *Endornaviridae* (Figure A4), except for its genome size of ∼21kb which is the longest genome reported to date for this group. In addition to the viral RdRp domain located in the Cter region of the Almopos endorna-like virus 1 protein, other protein domains and signatures could be identified that were related to the (+)RNA virus helicase core (IPR027351), the YbiA-like (IPR037238) and the UDP-Glycosyltransferase/glycogen phosphorylase superfamilies (SSF53756) (Figure A4), similar to the previous studies^43–46^. It is very likely that other viral proteins and functions are encoded but are too divergent to be identified. The investigation of these additional divergent viral translated products could be of significant importance for both revealing the evolutionary origins of the *Endornaviridae*^47^.

The Althepos endorna-like virus 1 sequence is only 4.8kb in length and likely represents a partial genome. Additional read mapping using our metagenomic or SRA-based data did not allow the retrieval of the full-length sequence (Figure A4). Whether Althepos endorna-like virus 1 and Almopos endorna-like virus 1 impact the fitness of their *G. carpenteri* host remains to be investigated but might have significant implications for the management of this potentially harmful species^48^. Considering the persistent lifestyle reported for endornaviruses^49^ and the high similarities in terms of RdRp homologies and genomic organisation, it is likely that Althepos endorna-like virus 1 and Almopos endorna-like virus 1 share the same infectious properties as other members of the genus *Alphaendornavirus* and might therefore constitute another example of capsid-less persistent viruses associated with protist hosts. Importantly, Althepos endorna-like virus 1 and Almopos endorna-like virus 1 identified from the *G. carpenteri* culture represent the second microalgal-endornavirus association observed to date^18^ and the first report in a dinoflagellate host, strongly suggesting that a microalgal-specific *Endornaviridae* clade may exist.

### A new Partitiviridae genus associated with dinoflagellate hosts

Orion durna-like virus 1 and Diktys durna-like virus 1, observed in *P. lima* (Dinophyceae) and *G. carpenteri* (Dinophyceae) cultures, respectively, form a clade with the Ourea durna-like virus previously associated with the dinoflagellate *Dinophysis acuminata*^20^ (Figure 5). Specifically, they form a sister clade to the genus *Deltapartitivirus*, belonging to the bi-segmented dsRNA *Partitiviridae* that infect fungi and plants^50^ and recently associated with unicellular algae^20^.

**Figure 5.**
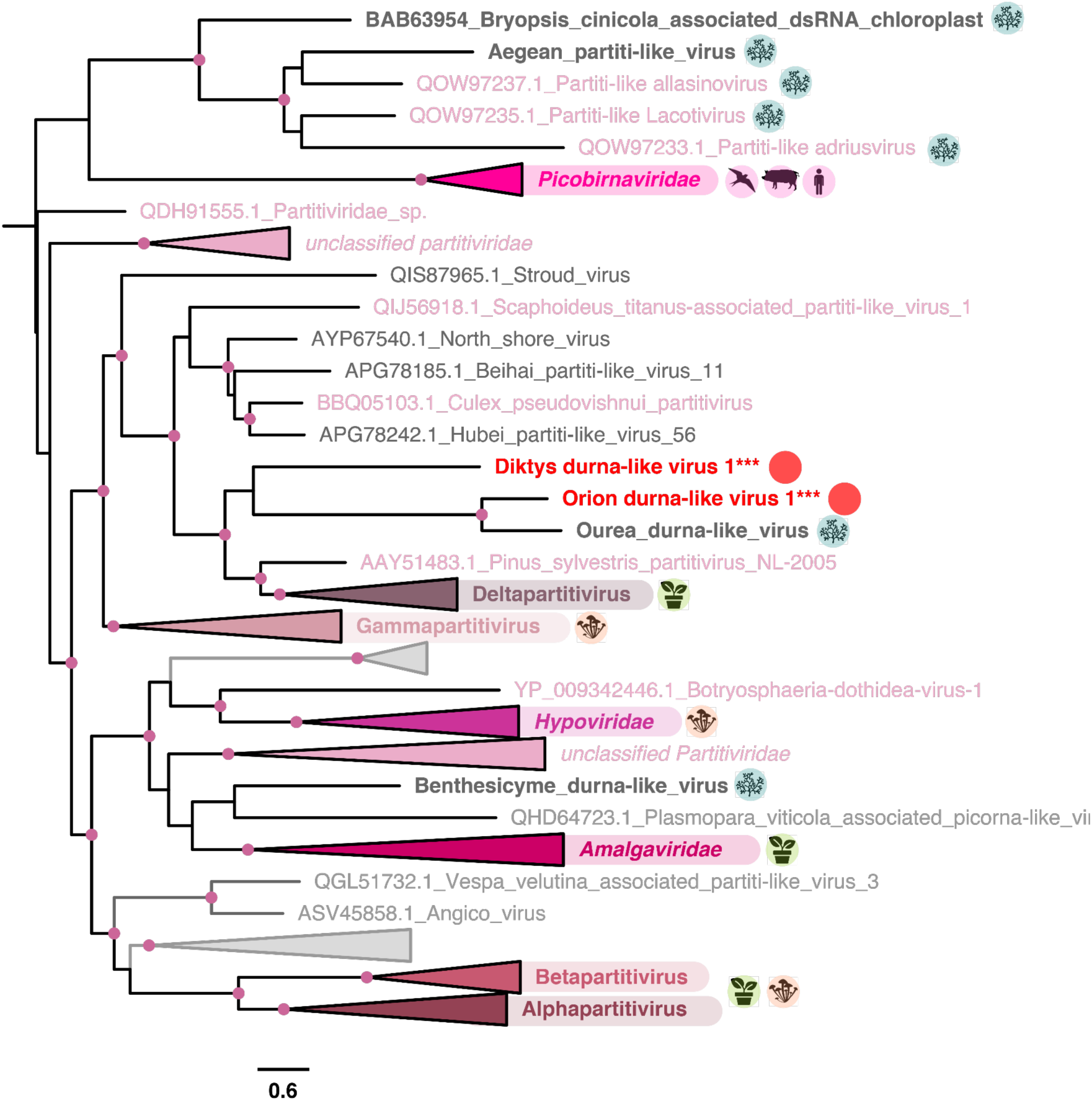
Phylogeny of the *Durnavirales*. Sequences in grey indicate unclassified viruses while those in bold refer to microalgal-associated viruses. Host lineages are indicated in circles to the right of major viral clades and correspond to metazoa (pink), fungi (orange), land plants (green), protozoa (light blue) and microalgae (blue). The new viral sequences identified in this study are indicated with a red circle. The tree is mid-rooted and confident nodes (with SH-alrt likelihood ratio test values >=80%) are represented as orange circles. The scale bar represents the number of amino acid substitutions per site.

Orion durna-like virus 1 and Diktys durna-like virus 1 genomes (1.8 kb and 2 kb in length, respectively) with a single ORF containing the RdRp domain (Figure A5). *Partitiviridae* are bisegmented viruses. Considering the placement of Orion durna-like virus 1 and Diktys durna-like virus 1 within the *Partitiviridae* phylogeny, it is very likely that they comprise a second segment, potentially encoding a coat protein not retrieved in this study due to our RdRp-based retrieval methodology. A complementary comparison of those sequences in the SRA database identified a sequence with 100% sequence identity at the amino acid level to Diktys durna-like virus 1 from a *Gambierdiscus polynesiensis* (dinoflagellate) sample (SRR3358210) (Table A3).

Together, these results are compatible with establishing a new *Partitiviridae* genus that is specific to dinoflagellates, comprising Orion durna-like virus 1, Diktys durna-like virus 1 and the previously identified Ourea durna-like virus. This observation expands the host range reported for this family, already comprising plants, fungi, oomycetes, apicomplexan parasites and green algae^19,51–54^. While most of the *Partitiviridae* do not induce symptoms in their hosts, hypovirulence has been reported in the alpha-, beta- and gammapartitiviruses^55^. Further analysis is required to assess the effects of Orion durna-like virus 1 and Diktys durna-like virus 1 on their potentially harmful dinoflagellate hosts *G. carpenteri* and *P. lima*^48,56^.

### *Marnaviridae*-like viruses associated with a *Nannochloropsis oculata* culture

Two newly identified viruses were identified in a *N. oculata* (a eustigmatophyte) sample and exhibited RdRp sequence similarity with the *Marnaviridae*, a picorna-like family of ss(+)RNA viruses that infect unicellular eukaryotes (Figure 6). With its taxonomy recently re-assessed to incorporate viruses from metagenomic studies^22^, the *Marnaviridae* are classified into seven genera. Accordingly, Minyas marna-like virus 1 belongs to the genus *Locarnavirus* that comprises viruses derived from marine environment, mollusc and fish-based metagenomic studies (Figure 6).

**Figure 6.**
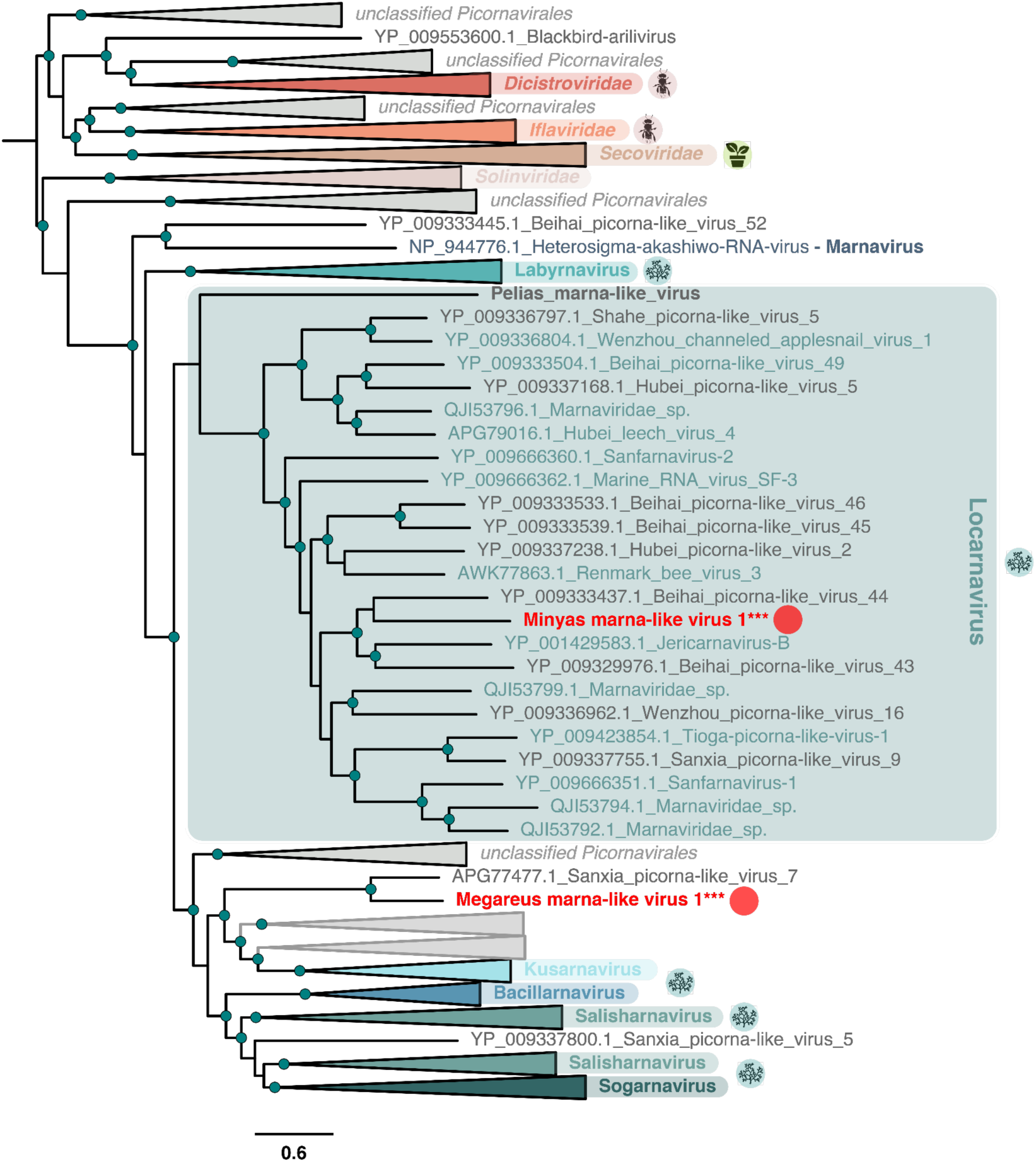
Phylogeny of the *Marnaviridae*. Sequences in grey indicate unclassified viruses while those in bold refer to algae-associated viruses. Host lineages are indicated in circles to the right of major viral clades and correspond to arthropods (pink), land plants (green), and microalgae (blue). The new viral sequences identified in this study are indicated with a red circle. The tree is mid-rooted and confident nodes (with SH-alrt likelihood ratio test values >=80%) are represented as orange circles. The scale bar depicts the number of amino acid substitutions per site.

This identification of Minyas marna-like virus 1 from the *N. oculata* culture provides compelling evidence for a *Locarnavirus* directly associated with a unicellular microalga. Along with the previous identification of the Dinophyceae-associated *Pelias marna-like virus*^20^ (Figure 6), this supports the idea of an extensive host range of locarnaviruses among unicellular microalgae. The second *Marnaviridae*-like virus, Megareus marna-like virus 1, forms a cluster with Sanxia picorna-like virus 7, falling in a position basal to Locarna-, Kusarna-, Bacillarna-, Salisharna- and Sogarnaviruses (Figure 6). It may therefore constitute a new genus of *Marnaviridae*^22^.

The short sequences of both Minyas marna-like virus 1 and Megareus marna-like virus 1 and their average read coverages (Table 1, Figure A5) strongly suggest that only partial genomes have been recovered. The low quantities of RNA extracted from the *N. oculata* sample and corresponding fragmented RNAs likely explain the poor coverage for the corresponding viral contigs, and that RT-PCR targeting both viral and host sequences returned negatives (Figure A2). Additional studies are needed to achieve the genomic and biological characterisation of those new *Marnaviridae*-like viruses associated with the eustigmatophyte host *N. oculata*. In particular, if the two newly reported *Marnaviridae* caused lysis of the biofuel-producing *N. oculata* cells this could represent a major concern for industrial-scale production.

### Divergent viruses identified using protein profiles and structural comparisons

To help identify viruses in basal and divergent microbial eukaryotes, we also conducted an approach based on HMM and structural RdRp comparisons, using the newly developed RdRp-scan tool^33^. Briefly, ORFs were predicted from each orphan contig and compared to RdRp profiles using Hidden Markov Models^33^. Such a strategy is expected to detect distant homologs sharing less than 30% of identity with viral protein sequences available in the current databases. As a result, two additional viral RdRp sequences were identified as distantly homologous to Pisuviricota members (Figure 2).

Using RdRp-scan HMM profiles a remote Pisuviricota-like RdRp signal was identified as associated with *E. gracilis*. The complementary Phyre2-based homology search returned a strong hit to the picornavirus sicinivirus 3dpol RdRp, thus validating the Pisuviricota-like signal previously detected. As noted above, comparison with *E. gracilis* nuclear (GCA_900893395) and mitochondrial (GCA_001638955) revealed strong identities (Table A2). A very close sequence (7 SNPs at the genome level and 6 non-synonymous substitutions at the RdRp level) could also be retrieved from the SRA database sample (SRR2294740), corresponding to the mitochondrial genome of *E. gracilis*. Such a mitochondrial sub-location is also suggested by the ORF found that can be expressed using the Chlorophycean mitochondrial genetic code (Figure A5). Hence, this Pisuviri-like signal appears to part of the host genome, likely corresponding to an endogenous viral element. Notably, the presence of such EVEs will help identify divergent viruses infecting euglenoid lineages, which are expected to be highly divergent considering the basal placement of euglenoids within eukaryotic organism diversity^32^.

The Phineus pisuviri-like virus 1, identified from *R. maculata*, was also confidently identified as a remote homolog of Pisuviricota RNA viruses using both RdRp-scan profiles and Phyre2 server. Although its distant and basal position in the RNA virus phylogeny prevents a robust comparison with existing *Riboviria* clades, its genome of 6.4kb and the associated read coverage suggests the full-length genome was recovered. The genome encodes three ORFs that possess RdRp function at the C-terminus (Figure A5). No functions could be associated with the additional ORFs and further studies are required to characterise this newly identified protist-infecting virus.

### Additional viruses

While it does not constitute a novel virus, one contig assembled from the *R. maculata* was retrieved in very high quantity and identical to the Despoena mito-like virus (Table 1), previously reported in another *R. maculata* sample^20^. This strongly reinforces the proposition of a Despoena mito-like virus infecting the red algal host and more generally the establishment of a mitovirus sub-clade that are able to infect microalgae^20^. Finally, our unbiased metagenomic analysis also retrieved additional viruses related to *Tombusviridae*, with identical sequences identified across several unrelated samples. It is very likely that these sequences result from contamination from kits or water used to extract or prepare RNA and cDNA libraries and were thus discarded from this study.

### Virus-Host assumptions based on the composition of microalgal cultures

The composition of the major kingdoms present in each sample was obtained by comparing contigs to the nt and nr database. The corresponding proportions as well as the abundance of contigs without a detectable match in nt or nr are shown in Figure 7. Bacteria-associated contigs are present within the libraries, especially those from *Nannochloropsis* and *Rhodella*, in line with the commonly reported microalgal-bacteria interactions^57^. Bacterial and eukaryotic viruses are usually too distantly related to be confounded. The presence of bacterial organisms in the samples is therefore not expected to interfere with our assumption that viruses identified as sharing homology with eukaryotic viruses very likely infect eukaryotic microalgal hosts. Remarkably, the proportion of undetected hits, without any match in nt and nr databases, is highly variable between libraries, ranging from less than 15% in the *T. weissflogii* sample to 40% in *G. carpenteri* culture (Figure 7A). This high variation likely arises from the lack of microalgal genomic and proteomic sequences in NCBI nt and nr databases, with genomic sequences available only for half of the microalgae hosts analysed here (Table A2). Such discrepancies in nucleotide and protein sequence assignment and abundance are further amplified in cases of highly abundant transcripts, such as ribosomal RNA, which very likely remain in the sample.

**Figure 7.**
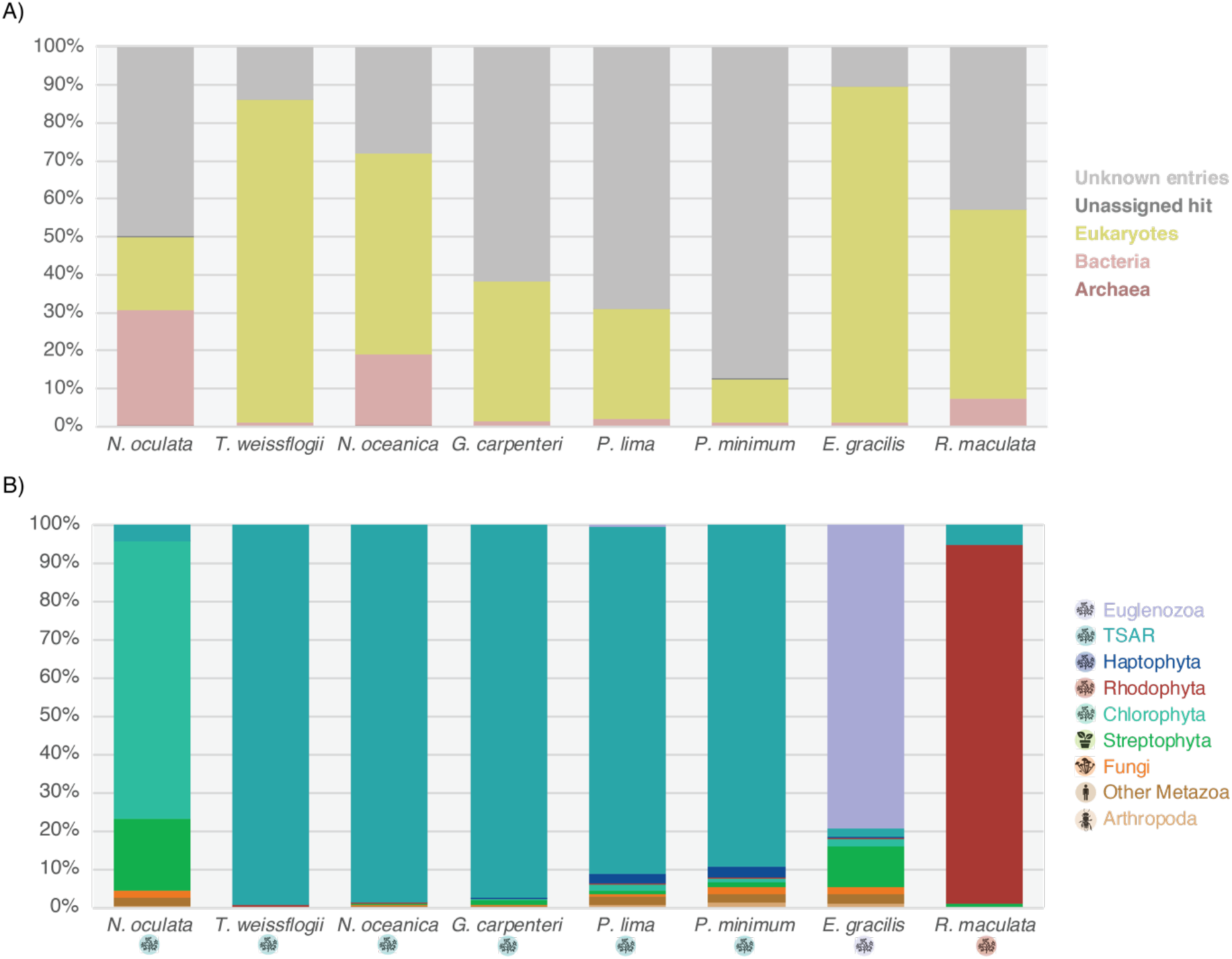
Relative abundance of contigs in microalgae libraries based on their assignment to major cellular organism clades. Contigs were assigned according to the taxonomy of their best Blast hits. Percentages of each contig were based on the abundance values and correspond to the sum of all contig TPM values belonging to each taxonomy clade. (A) Relative abundance of contigs associated with Kingdoms Archaea (dark pink), Bacteria (light pink) and Eukaryota (light yellow) using both BLASTn and BLASTx. The abundance of contigs with nt or nr entries lacking a taxonomy assignment are indicated in dark grey, while those without any nt or nr matches detected are indicated in light grey. (B) Relative abundances of contigs associated with major eukaryotic clades using BLASTx. Low abundance clades, counting for less than 0.5% of the total contig abundance, are not represented. TSAR: Telonemids-Stramenopiles-Alveolates-Rhizaria group as defined in ref.^32^.

While many unassigned entries in most of the samples analysed here can limit the formal assignment of viruses to hosts, obtaining a clearer picture of the eukaryotic host sequences present in the sample and their relative abundance can help to discriminate between eukaryotic hosts. Indeed, our cultures were washed several times before RNA extraction. It is therefore likely that the viral sequences identified result from intracellular viral forms rather than extracellular virions circulating in the culture media: hence, we assume that viruses detected in this study are associated with cellular organisms that are also present in the sample. We therefore examined the deep taxonomy of BLASTx eukaryotic-like contigs as well as the total contig abundance reported for major eukaryotic lineages (Figure 7B), which helped discriminate potential hosts for the most uncertain assignments. Accordingly, the very low abundance of fungi and land plant-associated sequences in *G. carpenteri* (Figure 7B) could constitute additional evidence for a microalgae-infecting endornavirus, even though members of the *Endornaviridae* have been traditionally associated to fungi or land plants (Streptophyta).

In the case of the Phineus pisuviri-like virus 1 identified from *R. maculata*, the majority of detectable contigs belong to the corresponding Rhodophyta host taxa, suggesting that this virus is likely associated with a Rhodophyte host rather than fungi or other contaminant organisms. The very large proportion of contigs associated with land plants (Streptophyta) in the *N. oculata* library might correspond to contamination. However, the unambiguous placement of the corresponding Minyas marna-like virus 1 virus within the well-established microalgae-infecting *Marnaviridae* provides a strong argument that this virus is associated with diatoms.

## Conclusions

Through metatranscriptomic sequencing of total RNA from microalgae cultures we identified ten new RNA viruses associated with diatom, eustigmatophyte, dinoflagellate and rhodophyte microalgae. These newly discovered viruses contribute to the establishment of new microalgae-infecting viral clades within the *Totiviridae* and *Partitiviridae*, as well as the enrichment of the positive single-stranded picorna-like family *Marnaviridae*. This study also extended the host range of the dsRNA *Endornaviruses* to microalgae, raising questions about how this viral family is able to infect the plant, fungi and TSAR eukaryotic supergroups. Considering the harmful or commercial value of their hosts, this description of new microalgal viruses paves the way for further studies of the effects of viral infections on host biology and their associated ecological and industrial consequences. Finally, this study highlights the need to reveal the hidden diversity among RNA viruses infecting microalgae, and to microbial eukaryotes in general, particularly considering their fundamental and applied importance.

## Materials and Methods

### Algae cultures

Microalgal cultures were maintained on a 12:12 light:dark cycle at 100 µmol m^-2^ s^-1^. Culture media and temperature conditions were specific to each species and were as follows: *Nannochloropsis oceanica* 24°C, f/2 medium; *Nannochloropsis oculata* 24°C, f/2 medium; *Thalassiosira weissflogii* 20°C, f/2 medium; *Rhodella maculata* 24°C, L1 medium (minus Si); *Euglena gracilis* 20°C, Euglena medium; *Prorocentrum* cf. *balticum* (UTSPH2D4)^48^; 20°C K medium-Si; *Prorocentrum lima* 25°C modified K medium^58^; *Gambierdiscus carpenteri* (UTSHI2C4) 25°C modified K medium^48,59^. To harvest each microalgal culture, the cells from 100-250 mL were pelleted by centrifugation at 200g for 4 mins and the supernatant discarded. The cells were then resuspended in 5 mL of artificial seawater and centrifuged again at 200g for 4 mins. This wash step was repeated twice more before a final centrifugation step at 1,000g for 4 mins followed by storage at -80°C until RNA extraction.

### Total RNA extraction and sequencing

Total RNA from the diatom (*T. weissflogii*) and Euglenozoa (*E. gracilis*) cultures were extracted using the RNeasy Plus Universal kit (Qiagen), according to the manufacturer’s instructions. Qiazol lysis buffer was then added to frozen pellets, and homogenisation was performed by pipetting. Genomic DNA was removed and RNAs extracted using 1-3 bromo-chloropropane. Supernatants were then transferred to Qiagen columns. After washing the columns, pure RNAs were collected into sterile water strictly following kit instructions.

Total RNA from dinoflagellates (*P. lima, P*. cf. *balticum, G. carpenteri*), the Rhodophyta *R. maculata* and the eustigmatophyte (*N. oceanica and N. oculata*) cultures was extracted using Allprep DNA/RNA kit (Qiagen), following the manufacturer instructions. Briefly, frozen cell pellets were supplanted with lysis RLT buffer and cells disrupted using bead beating with 0.5mm glass beads. An additional step of sample homogenisation using QIAshredder (Qiagen) was added during the RNA extraction of *R. maculata* sample and *N. oceanica* to reduce the viscosity of eluates. Cell debris was removed using a centrifugation step at high speed and the supernatants transferred to Qiagen columns. Total RNA fractions were then purified after several washing steps and eluted according to kit instructions.

### RNA sequencing

RNA quality was checked using a TapeStation and individually converted by the Australian Genome Research Facility (AGRF, Melbourne) into non-rRNA RNAseq libraries using TruSeq Stranded Total RNA with Ribo-Zero Plant (Illumina). Due to the very low RNA yields obtained for *Nannochloropsis oculata* and *Nannochloropsis oceanica*, these two libraries were prepared using the SMARTer Stranded Total RNA-Seq Kit v2—Pico Input Mammalian libraries (Takara Bio, Mountain View, CA, USA). The corresponding libraries were sequenced on the NovaSeq platform (Illumina) (paired-end, 150bp) by the AGRF.

### RNA-Seq data pre-processing: Read trimming, rRNA depletion and contig assembly

Total reads were filtered using Trimmomatic (v0.36)^60^ to remove low-quality and Illumina adapters. To maximize the completeness of the ribosomal (r) RNA depletion performed during library prep, the remaining rRNA reads were removed using the SortmeRNA program (2.1b)^61^. Filtered reads were then assembled into contigs using Trinity (v 2.5.1)^62^ and abundances (expected count and TPM) calculated using RSEM (v 1.3.1)^63^.

### Sample taxa composition

To help determine the taxa composition of each library, all contig sequences were compared to the non-redundant protein database nr from NCBI using Diamond BLASTx (v 2.0.9)^64^ and to the nucleotide database nt from NCBI using BLAST (v 2.2.30). The best hits were reported for each contig and their corresponding taxonomy analysed. For each library, contigs were grouped into major eukaryotic taxa and relative abundance determined as the sum of all the TPM (transcripts per million) within each taxon.

### RNA virus identification

#### Sequence-based similarity detection

RdRp sequences corresponding to RNA viruses (i.e. the *Riboviria*) were first identified by comparing contigs to the nr database using Diamond Blastx (v 2.0.9 ; e-value < 1e-05)^64^. To maximize the detection of RNA viruses, putative virus sequences identified from nr BLASTx as well as those previously obtained in an algae virus study^20^ were used as a database to perform a second round of BLASTx using contig libraries as queries and employing the same parameters as previously. The resulting RNA virus-like sequences were then submitted to the nr database (NCBI) and hits with the best match in cellular organism sequences were treated as false-positives and discarded from the analysis.

#### HMM-based homology detection of ORFans

All orphan contig sequences (i.e., that had no match in the nr database) were compared to the RdRp HMM-profiles of the RdRp-scan resource^33^ and using the HMMer3 program (v3.3)^65^.

#### Genome extension, Genome coverage and Virus annotation

To ensure all the RNA virus-like sequences could be identified and in their longest form, additional attempts to assemble contigs were performed using the rnaSPADES (v3.13.0)^66^ and Megahit programs (v1.2.9)^67^. This did not identify additional or longer RNA virus sequences. A manual elongation step was performed on viral candidates using Geneious (v11.1.4)^68^. A virus annotation to identify RdRp motifs was performed using InterProScan^69^ and RdRp-scan^33^. Genome coverage profiles were obtained by mapping the non-rRNA reads back to each contig sequence using Bowtie2 (v2.3.3.1)^70^ and Samtools (v1.6)^71^. The resulting SAM files were then plotted onto viral genomes using Geneious (v11.1.4)^68^.

#### SRA mining

To help retrieve complete genome sequences, assess intra-species variability and help associate viruses with particular algae hosts, we performed an additional step of Sequence Read Archive (SRA) mining for each of the ten new viruses identified in this study. For each algae library, we screened the SRA using nucleotide Magic-Blast (v1.3.0)^72^. When the number of hits exceeded 100, the corresponding SRA reads were mapped to the viral genome using Bowtie2 (v2.3.3.1)^70^ and SAMtools (v1.6)^71^.

#### Phylogenetic analysis

RNA virus phyla-level comparisons were performed using Clustal Omega (v1.2.4)^73^ to directly compare the newly identified sequences to the pre-built RdRp alignments from the RdRp-scan resource^33^. Initial phylogenetic trees were inferred using the maximum likelihood method available in FastTREE (v2.1.9; default parameters)^74^. Sub-alignments at the RNA virus order or family scale were then obtained using Clustal Omega (v1.2.4)^73^ and manually checked using Geneious (v11.1.4)^68^. Maximum likelihood phylogenies of these sub-alignments were then inferred using the IQ-TREE package (v2.0-rc1)^75^ with the best-fit amino acid substitution model obtained with ModelFinder Plus^76^ and using a Shimodaira-Hasegawa approximate-likelihood ratio and 1000 replicates (-alrt 1000) to assess nodal support.

### RT-PCR confirmation

To experimentally confirm viral contigs assembled from RNAseq data, cDNAs from each total RNAs were first obtained using the SuperScript IV reverse transcriptase (Invitrogen). PCRs were then performed on each cDNA sample using corresponding host and virus primers (detailed in Table A4) using the Platinum SuperFi II DNA polymerase (Invitrogen) and following manufacturer instructions.

## Data availability

Corresponding RNAseq read files will be available on the SRA under BioProject XXX, with accessions XXXX. Newly identified viral sequences will be deposited and available at GenBank/NCBI under the accessions XXXX.

## Acknowledgments

We thank Jean-Baptiste Raina for his useful comments on host compositions. ECH is supported by an Australian Research Council Australian Laureate Fellowship (FL170100022).

