## Supplementary Information for "Diverse RNA viruses associated with diatom, eustigmatophyte, dinoflagellate and rhodophyte microalgae cultures"

### Appendices

### Supplementary Tables

#### Table A1. Total RNA extractions and RNAseq results.

| **Algae species** | **Total RNA quantity** | **Sequencing data yields** |
|---|---|---|
| *Nannochloropsis oceanica* | 20ng | 27.81 Gb |
| *Nannochloropsis oculata* | 30ng | 52.54 Gb |
| *Thalassiosira weissflogii* | 105ng | 29.39 Gb |
| *Prorocentrum lima* | 700ng | 25.16 Gb |
| *Prorocentrum* cf. *balticum* | 7500ng | 40.80 Gb |
| *Gambierdiscus carpenteri* | 630ng | 51.80 Gb |
| *Rhodella maculata* | 120ng | 24.94 Gb |
| *Euglena gracilis* | 5700ng | 24.90 Gb |

**Table A2. Genome BLASTn analysis of the viruses detected.** Host algal species in which the closest species are used as surrogates are indicated with a * symbol.

| Viral Species | Algae Culture/Library | Algae Species | Genome | Blast |
|---|---|---|---|---|
| Triopas Ghabri-like Virus 1 | *P. cf. balticum* | *P. cf. balticum* | GCA_001652855.1 | No hits |
| Taphios Ghabri-like Virus 1 | *N. oceanica* | *N. oceanica* | GCA_004519485.1 | No hits |
| Taphios Ghabri-like Virus 1 | *T. weissflogii* | *T. oceanica** | GCA_019693575.1 | No hits |
| Diktys Durna-like Virus 1 | *P. lima* | *P. minimum** | GCA_001652855.1 | No hits |
| Orion Durna-like Virus 1 | *G. carpenteri* | *G. carpenteri* | Not available | ND |
| Almopos Endorna-like Virus 1 | *G. carpenteri* | *G. carpenteri* | Not available | ND |
| Althepos Endorna-like Virus 1 | *G. carpenteri* | *G. carpenteri* | Not available | ND |
| Phineus Pisuviri-like Virus 1 | *R. maculata* | *R. maculata* | Not available | ND |
| Megareus Marna-like Virus 1 | *N. oculata* | *N. oculata* | GCA_004335455.1 | No hits |
| Minyas Marna-like Virus 1 | *N. oculata* | *N. oculata* | GCA_004335455.1 | No hits |
| Pisuviri-like Signal | *E. gracilis* | *E. gracilis* | GCA_900893395.1<br>GCA_001638955<br>(mitochondrial) | +++ |

**Table A3. Number of SRA accessions screened for newly identified viruses and the corresponding hits.**

| Viral species | Library | Algae clade | SRA acc. nb | Hits found |
|---|---|---|---|---|
| Taphios ghabri-like virus 1 | *N. oceanica* | Nannochloropsis | 292 | None |
| Taphios ghabri-like virus 1 | *T. weissflogii* | Thalassiosira | 709 | SRR12347810 |
| Triopas ghabri-like virus 1.1 | *P. cf. balticum* | Prorocentrum | 143 | None |
| Diktys durna-like virus 1 | *P. lima* | Prorocentrum | 143 | None |
| Orion durna-like virus 1 | *G. carpenteri* | Gambierdiscus | 86 | SRR3358210 |
| Pisuviri-like signal | *E. gracilis* | Euglena | 72 | SRR2294740 |
| Almopos endorna-like virus 1 | *G. carpenteri* | Gambierdiscus | 86 | None |
| Althepos endorna-like virus 1 | *G. carpenteri* | Gambierdiscus | 86 | None |
| Phineus pisuviri-like virus 1 | *R. maculata* | Rhodella | 3 | None |
| Megareus marna-like virus 1 | *N. oculata* | Nannochloropsis | 292 | None |
| Minyas marna-like virus 1 | *N. oculata* | Nannochloropsis | 292 | None |

**Table A4. List of primer sequences used in this study.** All PCR reactions were performed at a universal annealing temperature of 60ºC. ITS: Internal Transcribed Spacer. LSU: Ribosomal Large Subunit.

| Primer ID | Direction | Sequence (5'-3') | Sequence targeted | length |
|---|---|---|---|---|
| Euglena-ITS Fw | Forward | TCCTGCCTATCACCCCACA | E. gracilis ITS | 273 |
| Euglena-ITS Rev | Reverse | CTACCCCGGTCCCGACTTT | | |
| Pisuviri-like signal Fw | Forward | TGCTGCACCTGCTATGCTT | Pisuviri-like signal | 500 |
| Pisuviri-like signal Rev | Reverse | CACGTGTGTCATCCCCACA | | |
| P.minimum_ITS Fw | Forward | CAGTTGGTGAGGCTCTGGG | P. *cf. balticum* ITS | 214 |
| P.minimum_ITS_Rev | Reverse | TCGTTGTTCGAGCCGAGAC | | |
| Triopas ghabri-like virus 1 | Forward | GCGARATGRTKGTSGARYT | Triopas ghabri-like virus 1 | 400 |
| Triopas ghabri-like virus 1 | Reverse | RTGSCCRATGWRCTGSACG | | |
| ITS1-Fw | Forward | TCCGTAGGTGAACCTGCGG | T.weissflogii ITS | 322 |
| Glaucocystis ITS_Rev | Reverse | TCGCTGCGTTCTTCATCGT | | |
| Taphios ghabri-like virus 1 | Forward | ACACGGAAAGGTGAGAGCG | Taphios ghabri-like virus 1 | 822 |
| Taphios ghabri-like virus 1 | Reverse | GGGCCAGCATAGCTAGCTC | | |
| N.oculata ITS | Forward | GTGGCCGATTATGGGAGGG | N. oculata ITS | 233 |
| Glaucocystis ITS | Reverse | TCGCTGCGTTCTTCATCGT | | |
| ITS1-FWD | Forward | TCCGTAGGTGAACCTGCGG | P. lima ITS | 403 |
| P. lima ITS | Reverse | TCAAGGGCCACAGCAAGAC | | |
| Diktys durna-like virus 1 | Forward | ACCGCATCTTCGCAACAGA | Diktys durna-like virus 1 | 664 |
| Diktys durna-like virus 1 | Reverse | AGCTTTGCATGCGGGTAGT | | |
| G_carpenteri_LSU | Forward | AGGCTGTGCATGGCTCATT | G. carpenteri LSU | 641 |
| G_carpenteri_LSU | Reverse | CAGCCATCCCCAGCAGAAA | | |
| Orion durna-like virus 1 | Forward | GCTCGGCTTCAGGTCAGTT | Orion durna-like virus 1 | 654 |
| Orion durna-like virus 1 | Reverse | CACTTGCGTTTTCGGTGGG | | |
| Althepos endorna-like virus 1 | Forward | AGTGGCGCACAAGCACTAT | Althepos endorna-like virus 1 | 877 |
| Althepos endorna-like virus 1 | Reverse | CCCCATTTCCAAGCCGTCT | | |
| Almopos endorna-like virus 1 | Forward | ACCTTCTTGGCCTGGATGC | Almopos endorna-like virus 1 | 623 |
| Almopos endorna-like virus 1 | Reverse | ATCCTCCTCAGACGGTGCT | | |
| Rhodella_maculata_ITS1 | Forward | CGGCCGAGTTCGACTATCC | R. maculata ITS1 | 759 |
| Rhodella_maculata_ITS1 | Reverse | TTCTTTGGGGTAGGCGCTG | | |
| Phineus pisuviri-like virus 1 | Forward | TCGGTCCAGATGGCAAACC | Phineus pisuviri-like virus 1 | 833 |
| Phineus pisuviri-like virus 1 | Reverse | CACCGGTGAGCCTTGTGAT | | |

### Supplementary Figures

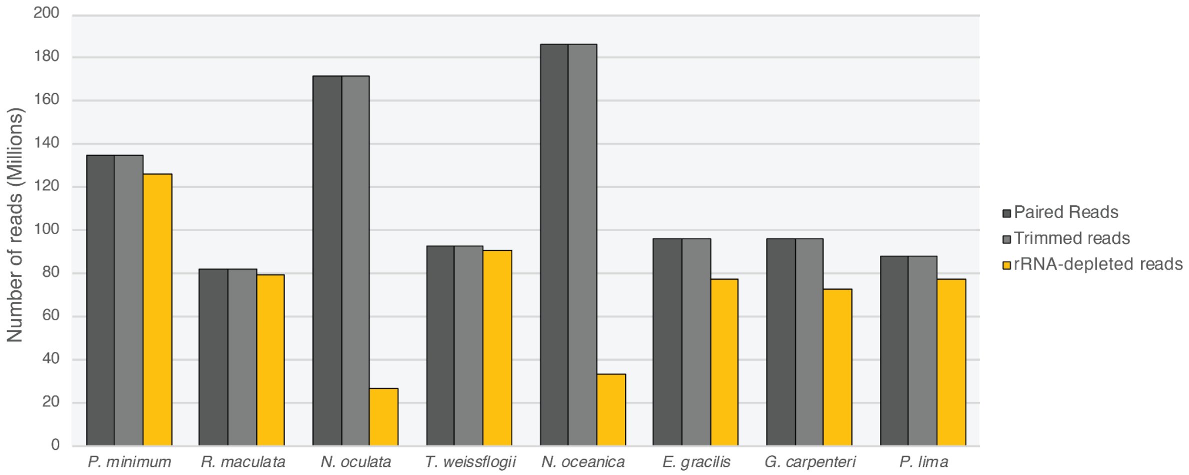


**Figure A1. Number of sequencing reads obtained for each algae library.** Paired-end reads obtained using Illumina NovaSeq high-throughput sequencing are indicated in dark grey. The final number of paired reads obtained after trimming and rRNA depletion steps are indicated in light grey and yellow, respectively.

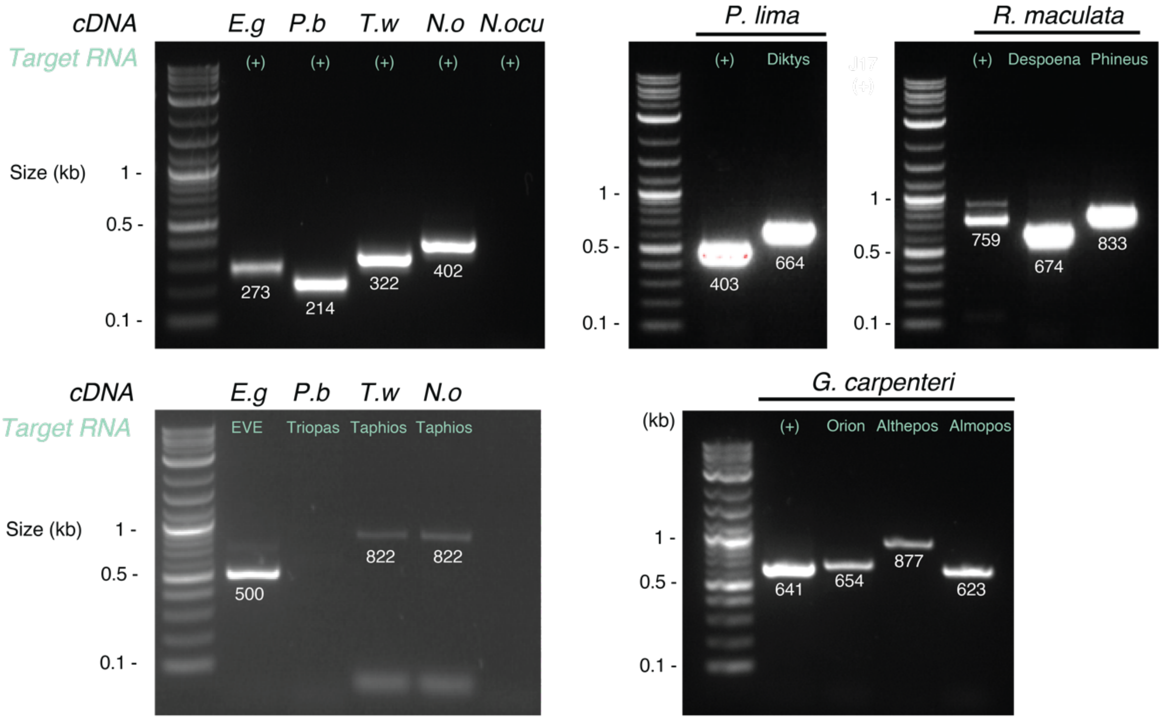


**Figure A2. RT-PCR confirmation of novel viral signals identified in this study.** The expected lengths of each PCR product are indicated below each band. RNA sequences targeted for each reaction are indicated in green. Corresponding RNA samples are indicated on top of each well (E.g: *Euglena gracilis*; P.b: *Prorocentrum cf. balticum*; T.w: *Thalassiosira weissflogii*; N.o: *Nannochloropsis oceanica*; N. ocu: *Nannochloropsis oculata*). (+): Host gene tested.

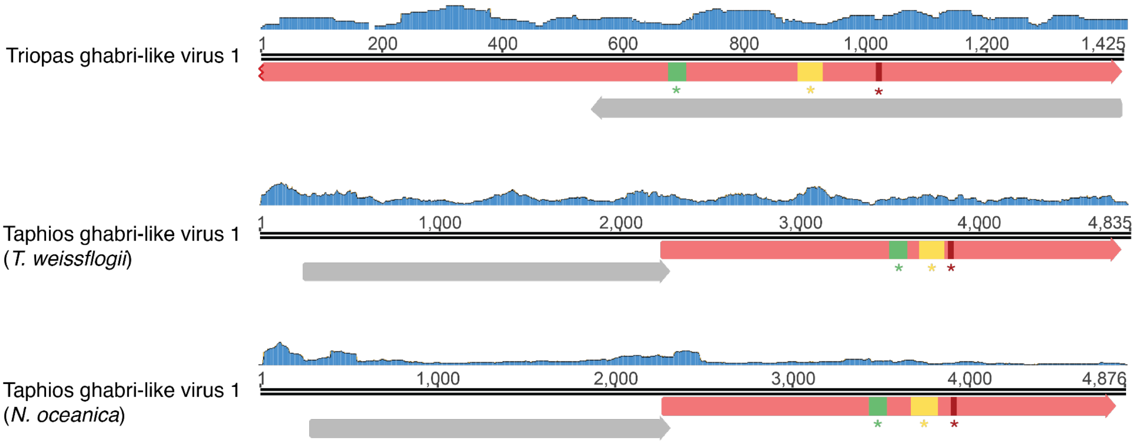


**Figure A3. Genome organisation of the Ghabri-like viruses identified in this study.** Read coverage of each genome is represented as a blue histogram. ORFs were predicted using standard genetic codes and their directions represented as arrows. ORFs encoding RdRp-like signals and hypothetical functions are indicated in red and grey, respectively. A, B and C RdRp motifs are indicated in green, yellow and red boxes, respectively.

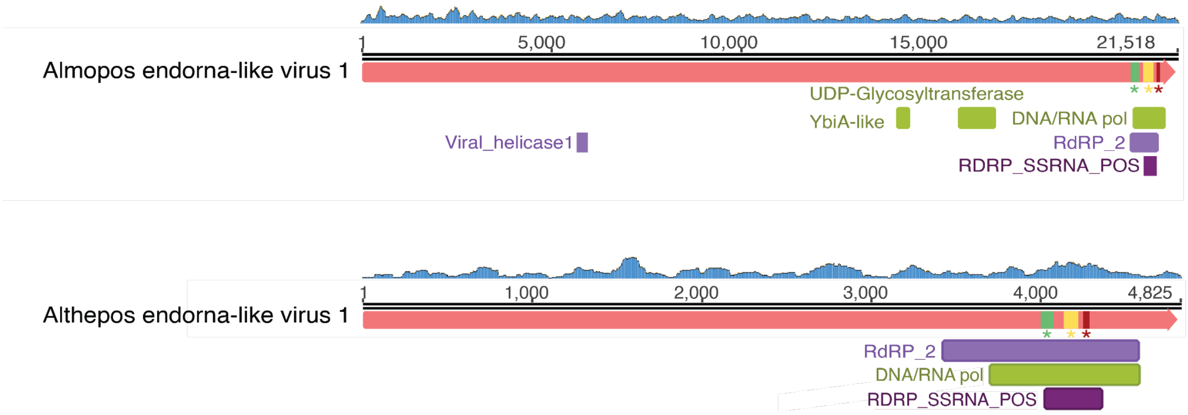


**Figure A4. Genome organisation of the Endorna-like viruses identified in this study.** Read coverage of each genome is represented as a blue histogram. ORFs were predicted using standard genetic codes and their directions represented as arrows. ORFs encoding RdRp-like signals and hypothetical functions are indicated in red and grey respectively. Light purple, green and dark purple boxes indicate PROSITE, SUPERFAMILY and PFAM predicted domains, respectively. A, B and C RdRp motifs are indicated in green, yellow and red boxes, respectively.

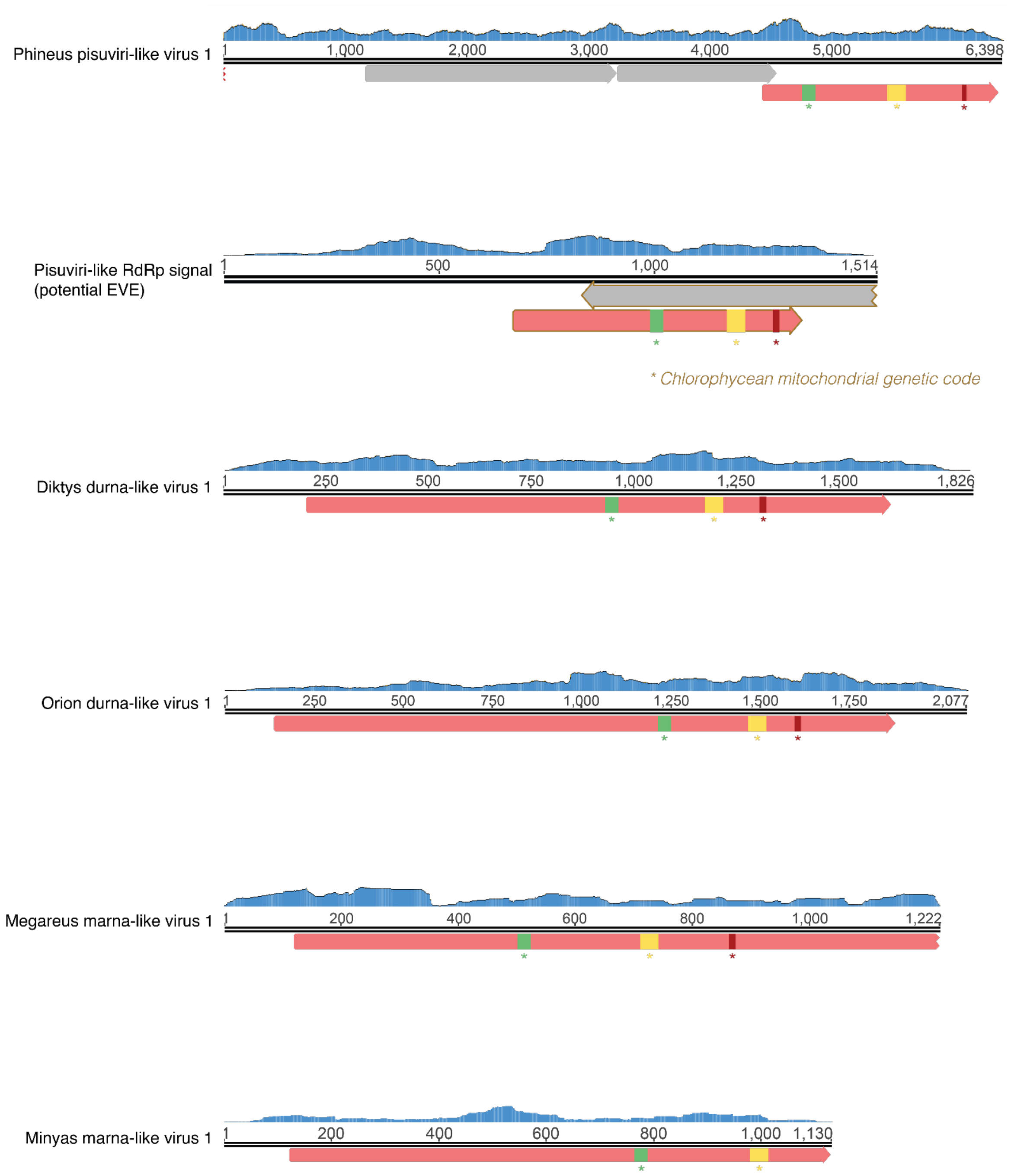


**Figure A5. Genome organisation of the Pisuviricota-like viral signals identified in this study.** Read coverage of each genome is represented as a blue histogram. ORFs were predicted using either standard genetic codes or Chlorophycean mitochondrial code, and their directions represented as arrows. ORFs encoding RdRp-like signals and hypothetical functions are indicated in red and grey, respectively. A, B and C RdRp motifs are indicated in green, yellow and red boxes, respectively.
